# The hepatic sinusoid is a site of microvascular complications of hyperglycemia

**DOI:** 10.64898/2026.10.15.759306

**Authors:** Cristina I. Øie, Gianina Dumitriu, Ruomei Li, Gahl Levy

## Abstract

Diabetes is the strongest independent predictor of progression of metabolic dysfunction-associated steatotic liver disease (MASLD) to cirrhosis and hepatocellular carcinoma, yet how glucose injures the liver is poorly understood. In the retina and kidney, hyperglycemia damages the microvascular endothelium through a shared mechanism in which unrestricted glucose entry inhibits glyceraldehyde-3-phosphate dehydrogenase (GAPDH) and diverts glycolytic intermediates into the four major pathways of glucotoxicity. The liver sinusoidal endothelial cell (LSEC) is the first cell exposed to portal blood, which carries glucose from the gut at concentrations higher than the systemic circulation reaches, for years before diabetes is diagnosed. LSEC capillarization, the loss of fenestrae and transition to a continuous capillary, precedes steatohepatitis and fibrosis. Here we show that the LSEC is a target of the same glucotoxic mechanism that underlies the classical diabetic complications. GLUT1 was the dominant glucose transporter of the LSEC. Hyperglycemia inhibited GAPDH in primary mouse LSECs and induced capillarization cell-autonomously, and inhibition of GAPDH alone reproduced the defenestration. Of the four glucotoxicity pathways, flux through the hexosamine pathway to protein O-GlcNAcylation was necessary for defenestration under high glucose and sufficient to induce it under low glucose, and was induced and inhibited in human LSECs. O-GlcNAcylation of endothelial nitric oxide synthase suppressed nitric oxide production, driving defenestration through cGMP and protein kinase G signaling and Rho kinase-driven actomyosin contraction. Intervening at any step of the cascade reversed the defenestration. Hyaluronan, cleared by the LSEC and normally catabolized through glycolysis, supplied a second, GFAT-independent input to protein O-GlcNAcylation under hyperglycemia. Loss of LSEC nitric oxide was sufficient to increase hepatocyte lipid accumulation in coculture. In mice with established diet-induced steatohepatitis, inhibiting O-GlcNAc transferase restored sinusoidal fenestrae and attenuated fibrosis. The hepatic sinusoid is therefore a site of microvascular complications of hyperglycemia, alongside the retina, kidney and peripheral nerve.

## Introduction

Metabolic dysfunction-associated steatotic liver disease (MASLD) affects more than a third of adults worldwide and approximately two of every three people with type 2 diabetes^1^. Diabetes is the strongest independent predictor of progression to cirrhosis and hepatocellular carcinoma^2,3^, and glycemic control improves fibrosis independent of body weight^4,5^. Hyperglycemia is therefore a candidate driver of MASLD progression rather than a passive comorbidity.

In the organs that develop diabetic complications, vascular endothelial cells are a major target of glucose-induced damage^6–8^. In these tissues glucose entry is not under insulin control, so intracellular glucose follows the blood concentration^6,7^. Under hyperglycemia, excess glucose uptake drives superoxide overproduction, which inhibits glyceraldehyde-3-phosphate dehydrogenase (GAPDH), and the upstream glycolytic intermediates accumulate and are shunted into the four major pathways of glucotoxicity^6–8^. These are augmented polyol flux, accumulation of advanced glycation end-product precursors, protein kinase C activation, and flux through the hexosamine biosynthetic pathway to protein O-GlcNAcylation^6,7^. The hepatic sinusoid has never been examined in this framework, although it is exposed to glucose at concentrations higher than the systemic circulation reaches, for years before diabetes is diagnosed^9–13^.

The liver sinusoidal endothelial cell (LSEC) is architecturally and functionally unlike any other endothelium in the body^14^. Its transcellular fenestrae, organized into sieve plates across a membrane devoid of a basement membrane, mediate size-selective exchange of lipoproteins and solutes between portal blood and hepatocytes^14,15^. Its scavenging receptor system, of exceptional capacity, clears hyaluronan, oxidized lipoproteins and macromolecular debris from portal blood^14,16–18^. Capillarization is the pathological dedifferentiation of this cell to a continuous capillary, defined by defenestration, deposition of a basement membrane and a shift from sinusoidal to continuous capillary identity^19,20^. Capillarization precedes steatohepatitis, fibrosis and hepatocyte ballooning^19–21^. Yet the molecular mechanism of capillarization remains unknown.

Glucose absorbed from the gut reaches the liver through the portal vein before entering the systemic circulation^9,10^. In insulin resistance, postprandial glucose excursions are greater^22–24^. LSECs line the hepatic sinusoids and are the first cells to contact portal blood^14^. We therefore hypothesized that portal hyperglycemia activates the four major pathways of glucotoxicity in the LSEC, and that flux through those pathways drives sinusoidal capillarization.

Here we identify the LSEC as a target of hyperglycemic damage and protein O-GlcNAcylation as a mechanism of sinusoidal capillarization. We show that high glucose capillarizes primary LSECs cell-autonomously and that GLUT1 is the dominant glucose transporter expressed by this cell. Metabolomic profiling showed that inhibition of GAPDH under low glucose raised the upstream glycolytic intermediates and lowered flux downstream, and the same inhibition phenocopied the defenestration caused by high glucose. Of the four pathways of glucotoxicity, inhibition of hexosamine flux significantly restored porosity under hyperglycemia. Raising O-GlcNAc under low glucose defenestrates the LSEC, and lowering O-GlcNAc under hyperglycemia increases porosity. We identify a second input to this pathway in the LSEC, in which hyaluronan scavenged by the LSEC raises protein O-GlcNAcylation, suggesting that salvaged GlcNAc is diverted into hexosamine flux downstream of glutamine:fructose-6-phosphate amido-transferase (GFAT) under hyperglycemia. Endothelial nitric oxide synthase (eNOS) is O-GlcNAcylated in the LSEC under hyperglycemia, and nitric oxide production falls. We establish the cascade from hyperglycemia to defenestration, through loss of nitric oxide, soluble guanylate cyclase, cGMP and protein kinase G to actomyosin contraction, by intervening at each step. Inhibiting nitric oxide synthesis in the LSEC raised hepatocyte lipid accumulation in coculture, and a nitric oxide donor lowered it. In mice with established diet-induced metabolic dysfunction-associated steatohepatitis (MASH), inhibition of protein O-GlcNAcylation restored sinusoidal fenestrations *in vivo*, attenuated fibrosis and improved serum markers of liver injury.

## Results

### Hyperglycemia inhibits GAPDH and capillarizes the LSEC

Capillarization is defined by defenestration, deposition of a basement membrane, and a shift from sinusoidal to continuous capillary identity^19,20^. To determine whether hyperglycemia induces capillarization in the LSEC, we cultured freshly isolated primary mouse LSECs for 24 h in 5.5 mM (low) or 30 mM (high) glucose. LSEC porosity, the percentage of cell area covered in fenestrations, fell by 67% under high glucose (P = 0.0029; n = 3) (Fig. 1a). Mannitol, used as an osmotic control, had no effect on porosity (P = 0.87; n = 3) (Supplementary Fig. 1). High glucose reduced expression of the LSEC-specific genes *Stab1, Stab2* and *Fcgr2b* by 17%, 24% and 24% (P = 0.034, P = 0.014, P = 0.008, respectively; n = 5), with a reciprocal 29% increase in the continuous endothelial marker *Cd34* (P = 0.042; n = 5), whereas *Pecam1* rose 17% without reaching significance (P = 0.26; n = 5) (Fig. 1b). Hyperglycemia is therefore sufficient to capillarize the LSEC cell-autonomously.

**Fig. 1.**
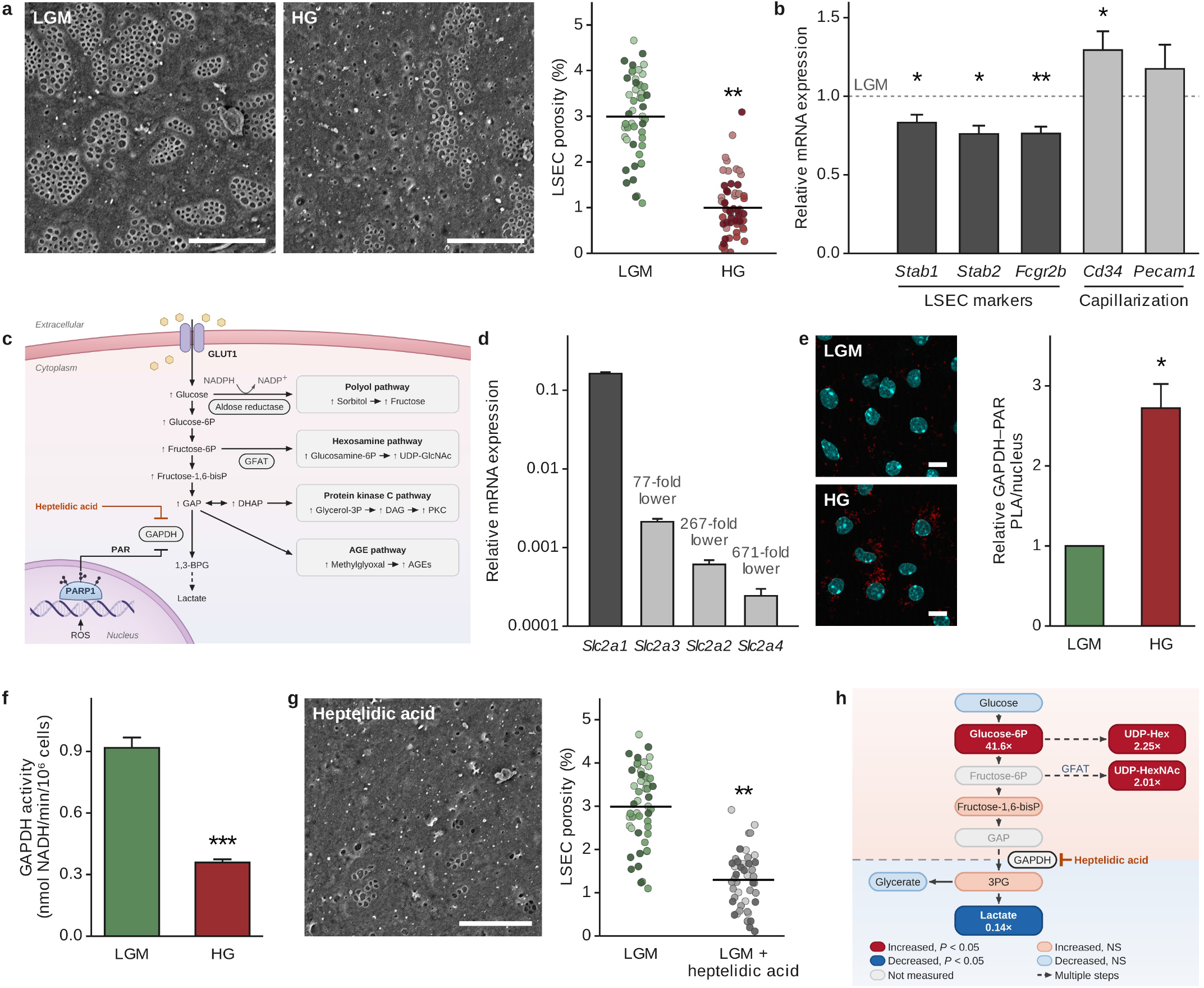
High glucose inhibits GAPDH and capillarizes the LSEC. Primary mouse LSECs were cultured for 24 h in low glucose with mannitol (LGM, 5.5 mM) or high glucose (HG, 30 mM). **a,** High glucose decreases LSEC porosity, the percentage of cell surface occupied by fenestrae. Representative scanning electron micrographs and quantification. Each dot is one cell, shaded by preparation. Horizontal line, mean of preparation means (n = 3 preparations, two-tailed unpaired t-test). **b,** High glucose decreases the LSEC identity genes Stab1, Stab2 and Fcgr2b and increases the capillarization marker Cd34, relative to LGM, dashed line (n = 5 preparations, one-sample t-test on log2 ratios). **c,** Glycolysis and the four glucotoxicity pathways that branch above GAPDH. Based on refs. 6,7. **d,** Slc2a1 (GLUT1) is the dominant glucose transporter of the LSEC. Transcripts relative to reference genes, with fold differences from Slc2a1 above the bars (n = 15 to 20 samples from 5 preparations). **e,** High glucose increases poly-ADP-ribosylation of GAPDH. GAPDH–PAR proximity ligation signal per nucleus, relative to LGM (n = 3 preparations, one-sample t-test on log2 ratios). **f,** High glucose decreases GAPDH activity (n = 3 preparations, two-tailed unpaired t-test). **g,** Inhibition of GAPDH with heptelidic acid decreases LSEC porosity under low glucose. Dots and line as in a (n = 3 preparations, two-tailed unpaired t-test). **h,** Heptelidic acid increases glucose 6-phosphate and UDP-HexNAc and decreases lactate. Fold change against control, color as keyed. UDP-Hex, UDP-glucose and UDP-galactose combined. UDP-HexNAc, UDP-GlcNAc and UDP-GalNAc combined (n = 5 preparations). Error bars, s.e.m. *P *<* 0.05, **P *<* 0.01, ***P *<* 0.001 versus LGM. Scale bars, 3 µm (a, g) and 10 µm (e).

Diabetic complications arise in cells that cannot limit glucose entry^6,25^. In vascular endothelial cells, glucose enters through GLUT1 independently of insulin^25^. Under hyperglycemia, superoxide causes DNA strand breaks that activate poly-ADP-ribose polymerase (PARP), which ribosylates and inactivates GAPDH, diverting the upstream glycolytic intermediates into the four major pathways of glucotoxic-ity (Fig. 1c)^6,7^. We asked whether capillarization of the LSEC arises through this same mechanism, beginning with how glucose enters the cell. *Slc2a1*, encoding GLUT1, was dominant, exceeding every other plasma-membrane transporter by at least 77-fold. *Slc2a4*, encoding the insulin-responsive GLUT4, was 671-fold lower (Fig. 1d). *Slc2a1* expression was not downregulated by high glucose, rising 17% (P = 0.038; n = 5) (Supplementary Fig. 2). Intracellular glucose in the LSEC is therefore set by the concentration in portal blood.

We then tested whether GAPDH is inhibited in the LSEC under hyperglycemia. Poly-ADP-ribosylation of GAPDH rose 2.72-fold as measured by proximity ligation (P = 0.011; n = 3) (Fig. 1e), while GAPDH activity fell by 61% (P = 0.0004; n = 3) (Fig. 1f). We therefore inhibited GAPDH under low glucose to test whether loss of its activity alone is sufficient to capillarize the LSEC. Heptelidic acid, an inhibitor of GAPDH, reduced porosity by 57% under low glucose (P = 0.0018; n= 3) (Fig. 1g). Metabolomic profiling of LSECs treated with heptelidic acid under low glucose confirmed the accumulation of intermediates upstream of GAPDH. Glucose 6-phosphate rose 41.5-fold (P *<* 0.0001; n = 5), lactate fell by 86% (P = 0.00003; n = 5), and UDP-HexNAc rose 2.0-fold (P = 0.0007; n = 5) (Fig. 1h, Supplementary Table 1).

The LSEC takes up glucose through the same insulin-independent transporter as the classical diabetic target tissues, sustains the same inhibition of GAPDH, and reproduces the loss of fenestrae when GAPDH is inhibited under low glucose. These results place the hepatic sinusoid among the sites of microvascular complications of hyperglycemia.

### Hexosamine flux to protein O-GlcNAcylation capillarizes the LSEC

To determine which of the four major pathways of glucotoxicity are required for the loss of fenestrae, we inhibited each pathway individually under hyperglycemia and measured porosity (Fig. 1c, 2a). Only inhibition of hexosamine flux with 6-diazo-5-oxo-L-norleucine (DON) significantly increased porosity, to 96% of the low glucose level (P = 0.038; n = 3). Inhibition of aldose reductase with zopolrestat, of advanced glycation with pyridoxamine, and of protein kinase C β with ruboxistaurin increased it to 87%, 73% and 50% without reaching significance (P = 0.077, P = 0.22 and P = 0.83, respectively; n = 3).

**Fig. 2.**
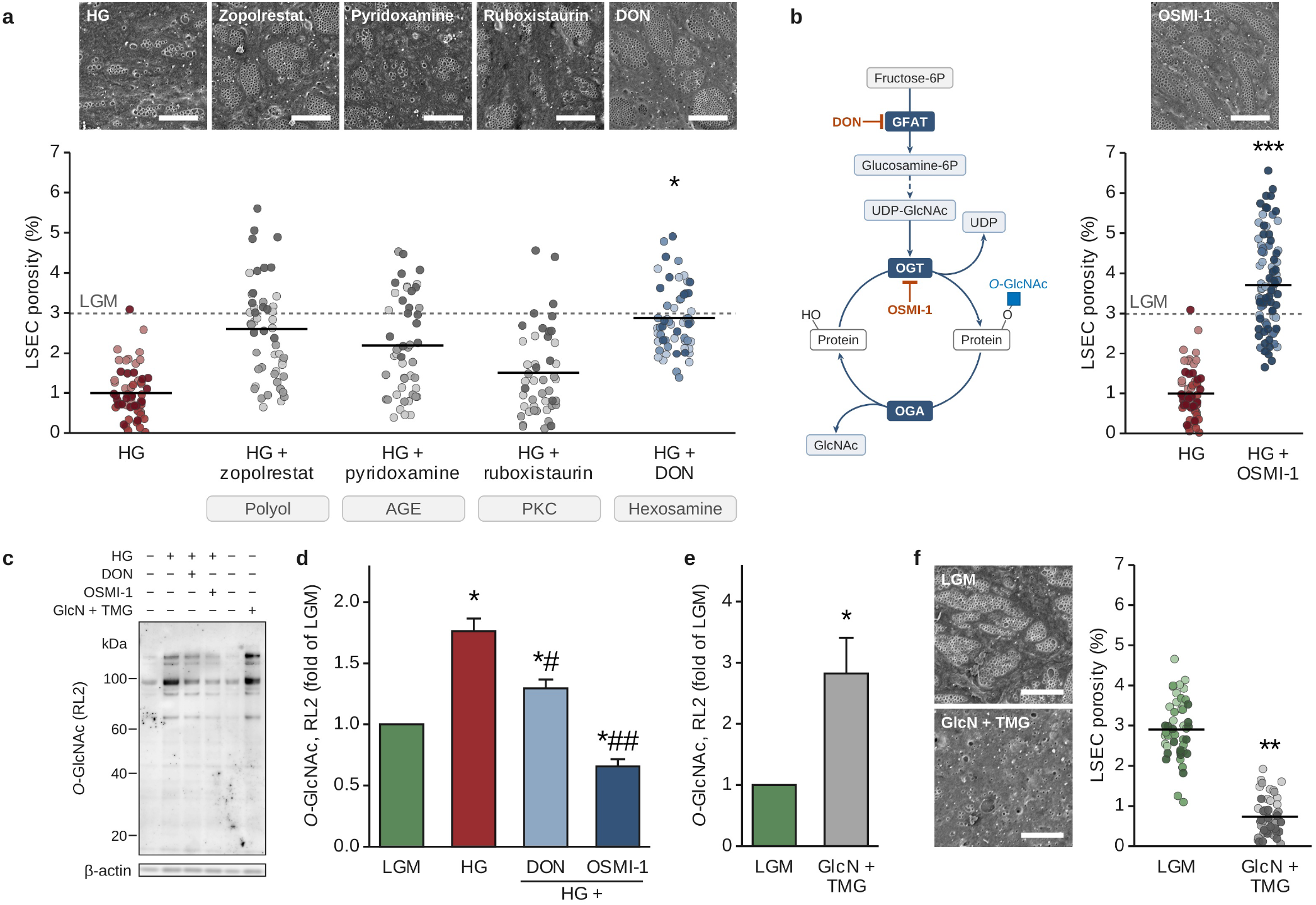
Hexosamine flux to protein O-GlcNAcylation is necessary and sufficient for defenestration. a, Inhibition of GFAT with DON prevents the loss of LSEC porosity in HG. Inhibition of the other three pathways with zopolrestat (aldose reductase), pyridoxamine (advanced glycation) or ruboxistaurin (protein kinase C) does not significantly increase LSEC porosity (n = 3 preparations, one-way ANOVA with Dunnett’s test). **b,** The hexosamine pathway and O-GlcNAc cycle. Inhibition of OGT with OSMI-1 prevents the loss of LSEC porosity in HG (n = 3 preparations, two-tailed unpaired t-test). **c,** Representative O-GlcNAc (RL2) immunoblot. **d,** High glucose increases global protein O-GlcNAcylation, and DON and OSMI-1 decrease it. Whole-lane RL2 signal, normalized to β-actin and relative to LGM (n = 3 preparations, one-sample t-test on log2 ratios). **e,** Glucosamine and the O-GlcNAcase inhibitor Thiamet-G (GlcN + TMG) increase global protein O-GlcNAcylation under low glucose. Whole-lane RL2 signal, normalized to β-actin and relative to LGM (n = 3 preparations, one-sample t-test on log2 ratios). **f,** GlcN + TMG decreases LSEC porosity under low glucose. Dots and line as in Fig. 1a. Dashed line, LGM mean (n = 3 preparations, two-tailed unpaired t-test). Error bars, s.e.m. *P *<* 0.05, **P *<* 0.01, ***P *<* 0.001 versus HG (a, b) or LGM (d–f). #P *<* 0.05, ##P *<* 0.01 versus HG. Scale bars, 3 µm.

DON inhibits GFAT, the rate-limiting reaction that commits fructose-6-phosphate to the hexosamine biosynthetic pathway^26^. The pathway produces uridine diphosphate N-acetylglucosamine (UDP-GlcNAc), the donor for protein O-GlcNAcylation, which is catalyzed by O-GlcNAc transferase (OGT) on serine and threonine residues^27–29^. OSMI-1, an inhibitor of OGT^30^, restored porosity to 124% of the low glucose level (P = 0.0004; n = 3) (Fig. 2b).

We measured global protein O-GlcNAcylation by RL2 immunoblotting (Fig. 2c). Hyperglycemia increased the signal to 176% of the low glucose level (P = 0.010; n = 3) (Fig. 2d). Inhibition of GFAT with DON and of OGT with OSMI-1 lowered it to 129% and 66% of the low glucose level (P = 0.043 and P = 0.002, respectively).

To test whether protein O-GlcNAcylation alone is sufficient to capillarize the LSEC, we treated LSECs under low glucose with glucosamine, which is phosphorylated to glucosamine 6-phosphate and bypasses the rate-limiting reaction catalyzed by GFAT^26^. Thiamet-G, an inhibitor of O-GlcNAcase^31,32^, was added to prevent removal of the modification^33^ (Fig. 2c). Global protein O-GlcNAcylation rose 2.8-fold (P = 0.032; n = 3) (Fig. 2e) and porosity fell to 25% of the low glucose level (P = 0.0019; n = 3) (Fig. 2f). We then treated primary human LSECs with glucosamine and Thiamet-G. Protein O-GlcNAcylation rose, and OSMI-1 lowered the modification (Supplementary Fig. 3).

Flux through the hexosamine biosynthetic pathway to protein O-GlcNAcylation is therefore necessary for capillarization under hyperglycemia and sufficient to capillarize the LSEC under low glucose.

### Loss of nitric oxide signaling defenestrates the LSEC

Nitric oxide generated by eNOS activates soluble guanylate cyclase (sGC) to convert GTP to cGMP, and cGMP activates protein kinase G (PKG)^34^. PKG phosphorylates vasodilator-stimulated phosphoprotein (VASP), which limits actin polymerization^35^. PKG also phosphorylates RhoA, which suppresses Rho kinase (ROCK) and reduces myosin light chain phosphorylation^36^, lowering cortical cytoskeletal tension. In LSECs, inhibition of ROCK or depolymerization of actin increases fenestrations^37–40^. What sets this cascade in motion during capillarization is not known.

eNOS is a substrate of OGT, and its O-GlcNAcylation in endothelial cells has been implicated in the microvascular and macrovascular complications of diabetes^41–44^. We therefore hypothesized that O-GlcNAcylation of eNOS in the LSEC lowers nitric oxide production, and that the resulting loss of cGMP and PKG activity increases actin polymerization and actomyosin tension, driving defenestration (Fig. 3a).

**Fig. 3.**
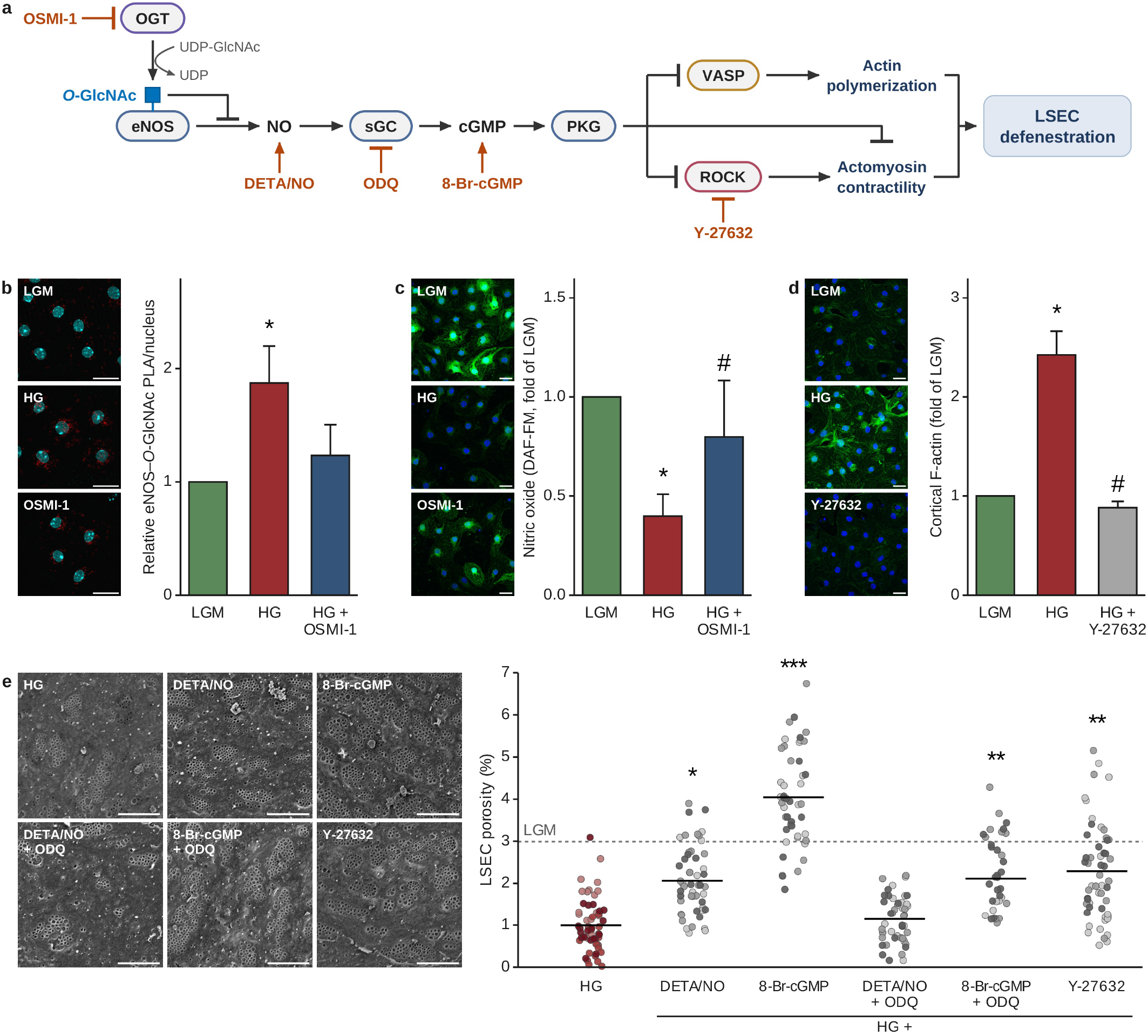
High glucose O-GlcNAcylates eNOS and defenestrates the LSEC through loss of nitric oxide signaling. a, Proposed pathway from eNOS O-GlcNAcylation to defenestration, with the agents used to test each step. **b,** High glucose increases O-GlcNAcylation of eNOS, and OSMI-1 returns it to the low glucose level. eNOS–O-GlcNAc proximity ligation, relative to LGM (n = 4 preparations, one-sample t-test on log2 ratios). **c,** High glucose decreases nitric oxide production, and OSMI-1 prevents the decrease. DAF-FM fluorescence, relative to LGM (n = 4 preparations, one-sample t-test on log2 ratios). **d,** High glucose increases cortical F-actin, and the ROCK inhibitor Y-27632 decreases it. F-actin intensity, relative to LGM (n = 3 preparations, one-sample t-test on log2 ratios). **e,** The nitric oxide donor DETA-NONOate (DETA/NO), the cGMP analog 8-Br-cGMP and Y-27632 each prevent the loss of LSEC porosity in HG. The soluble guanylate cyclase inhibitor ODQ blocks the effect of DETA/NO but not of 8-Br-cGMP. Dots and line as in Fig. 1a. Dashed line, LGM mean (n = 3 preparations, one-way ANOVA with Dunnett’s test). Error bars, s.e.m. *P *<* 0.05, **P *<* 0.01, ***P *<* 0.001 versus LGM (b–d) or HG (e). #P *<* 0.05, ###P *<* 0.001 versus HG. Scale bars, 20 µm (b–d) and 3 µm (e).

To test whether eNOS is O-GlcNAcylated in the LSEC, we used proximity ligation between eNOS and the O-GlcNAc antibody RL2 (Fig. 3b). Hyperglycemia increased O-GlcNAcylation of eNOS to 187% of the low glucose level (P = 0.030; n = 4), and OSMI-1 returned it to 123%, which was not significantly different from low glucose (P = 0.37; n = 4). Nitric oxide production fell to 40% of the low glucose level (P = 0.032; n = 4), and OSMI-1 increased it 2.0-fold (P = 0.015; n = 4), to 80% of the low glucose level (P = 0.51) (Fig. 3c). *Nos3* expression was unchanged by high glucose (97% of the low glucose level, P = 0.75; n = 5) (Supplementary Fig. 4), so the loss of nitric oxide does not reflect lower eNOS expression. Hyperglycemia therefore increases O-GlcNAcylation of eNOS and lowers nitric oxide output, and inhibition of protein O-GlcNAcylation reverses both.

The loss of nitric oxide should lower PKG activity and promote actin polymerization. ROCK activation reorganizes cortical actin into contractile bundles^45^. Under hyperglycemia actin intensity rose 2.4-fold (P = 0.011; n = 3), and the ROCK inhibitor Y-27632 lowered it to 36% of the hyperglycemic level (P = 0.023; n = 3), returning it to 88% of the low glucose level (P = 0.21) (Fig. 3d).

We next asked whether intervening at each step of this cascade restores porosity in hyperglycemic LSECs (Fig. 3e). The nitric oxide donor DETA-NONOate raised porosity 2.1-fold (P = 0.010; n = 3). The membrane-permeable cGMP analog 8-Br-cGMP raised porosity 4.1-fold (P *<* 0.0001; n = 3), and the ROCK inhibitor Y-27632 2.3-fold (P = 0.0024; n = 3). We then blocked sGC with ODQ, which acts down-stream of nitric oxide and upstream of cGMP (Fig. 3a). ODQ abolished the rescue by DETA-NONOate, leaving porosity at 115% of the hyperglycemic level (P = 0.97; n = 3), but not the rescue by 8-Br-cGMP, which still raised porosity 2.1-fold (P = 0.007; n = 3) (Fig. 3e). The effect of nitric oxide on porosity is therefore cGMP-dependent.

These results indicate that hyperglycemia O-GlcNAcylates eNOS and suppresses nitric oxide/sGC/cGMP/PKG signaling, releasing ROCK, which increases the actin polymerization and actomyosin contraction that defenestrate the LSEC.

### Hyaluronan salvage is a second input to hexosamine flux in the LSEC

The LSEC is the principal site of clearance of circulating hyaluronan, which it takes up and degrades in lysosomes to free N-acetylglucosamine (GlcNAc) and glucuronic acid, which are subsequently exported to the cytosol^46,47^. GlcNAc is degraded to acetate and lactate in these cells^46^, so it is metabolized through glycolysis.

Free GlcNAc is not a substrate for hexokinase, so entry into glycolysis has to occur at fructose-6-phosphate. Reaching fructose-6-phosphate requires three consecutive reactions, phosphorylation at C6, deacetylation and deamination, each catalyzed by a single known mammalian enzyme family (Fig. 4a). The deacetylation has been demonstrated in the LSEC^48^, where the acetate released is recovered from degraded hyaluronan^46^. We therefore reasoned that the LSEC disposes of salvaged GlcNAc through N-acetylglucosamine kinase (NAGK), the N-acetylglucosamine-6-phosphate deacetylase AMDHD2 and the glucosamine-6-phosphate deaminase GNPDA1 or GNPDA2. Expression of these enzymes in the LSEC has not been reported. All four were expressed, most abundantly the deaminase GNPDA2 that catalyzes the final step to fructose-6-phosphate and entry into glycolysis (Fig. 4b).

**Fig. 4.**
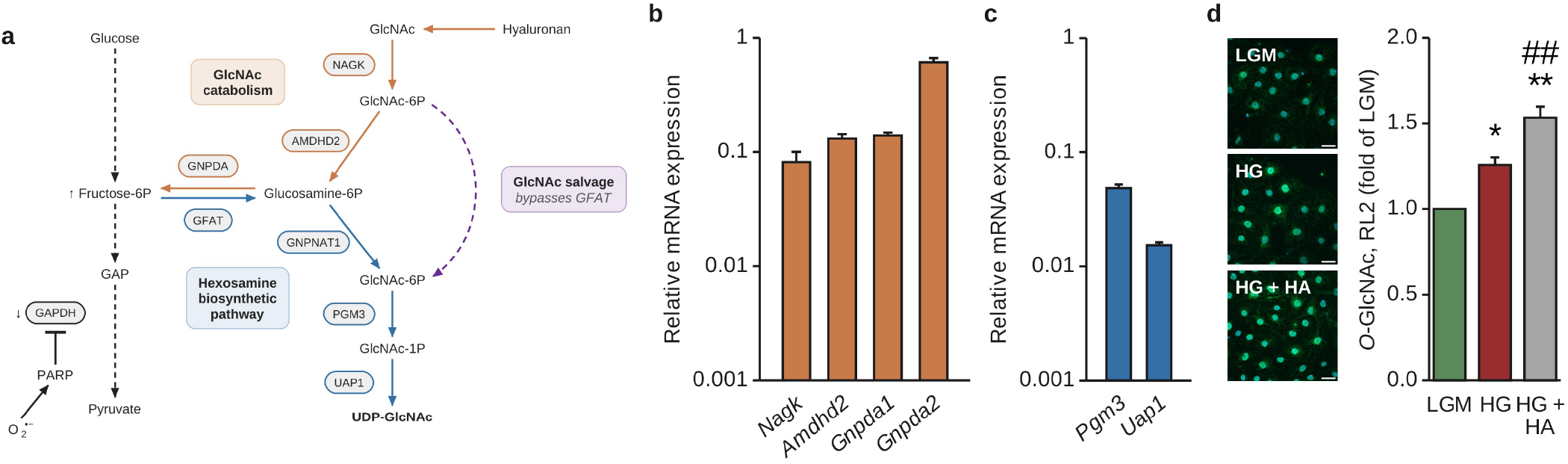
Hyaluronan supplies a GFAT-independent input to O-GlcNAcylation in the LSEC. a, Proposed diversion of hyaluronan-derived GlcNAc. GlcNAc enters as GlcNAc-6-phosphate, downstream of GFAT, and is normally catabolized to fructose-6-phosphate. Under high glucose it is diverted to UDP-GlcNAc. **b,** The LSEC expresses the enzymes of the catabolic route, Nagk, Amdhd2, Gnpda1 and Gnpda2. Transcripts relative to reference genes (n = 20 samples from 5 preparations). **c,** The LSEC expresses Pgm3 and Uap1, the enzymes of the route to UDP-GlcNAc. Transcripts relative to reference genes, on the same scale as b (n = 20 samples from 5 preparations). **d,** High glucose increases protein O-GlcNAcylation, and hyaluronan increases it further. RL2 immunofluorescence, relative to LGM (n = 3 preparations, one-sample t-test on log2 ratios). Error bars, s.e.m. *P *<* 0.05, **P *<* 0.01 versus LGM. ##P *<* 0.01 versus HG. Scale bars, 20 µm.

GlcNAc-6-phosphate, the first intermediate of the salvage route, is also an intermediate of hexosamine biosynthesis, downstream of the rate-limiting step catalyzed by GFAT (Fig. 4a). At that node, two enzymes compete for GlcNAc-6-phosphate. AMDHD2 directs it toward glycolysis and the mutase PGM3 toward UDP-GlcNAc, the donor for protein O-GlcNAcylation^27,49,50^. The deaminase reaction that produces fructose-6-phosphate is freely reversible^51^, so its net direction is set by the fructose-6-phosphate concentration in the cell. We therefore hypothesized that when fructose-6-phosphate accumulates under hyperglycemia the glycolytic disposal route at the deaminase is no longer favored, and GlcNAc-6-phosphate is diverted toward UDP-GlcNAc, raising protein O-GlcNAcylation. PGM3 and UAP1 were both expressed in the LSEC (Fig. 4c). The LSEC also expressed the GFAT genes *Gfpt1* and *Gfpt2* and the O-GlcNAc cycling enzymes *Ogt* and *Oga* (Supplementary Fig. 5). We supplied hyaluronan to LSECs under high glucose and measured protein O-GlcNAcylation by RL2 immunofluorescence. Hyaluronan raised protein O-GlcNAcylation to 122% of the high glucose level (P = 0.0011; n = 3) (Fig. 4d). Free HexNAc also rose in the metabolome when GAPDH was inhibited, by 53%, although this did not survive correction for multiple testing (Supplementary Table 1).

Under hyperglycemia, hyaluronan clearance therefore feeds the hexosamine flux that capillarizes the LSEC.

### Hyperglycemia in the LSEC drives hepatocyte lipid accumulation

Nitric oxide activates AMP-activated protein kinase^52^, and the resulting inhibitory phosphorylation of acetyl-CoA carboxylase suppresses de novo lipogenesis in the hepatocyte^53–55^. Hyperglycemia lowers nitric oxide production in the LSEC (Fig. 3c). We therefore tested whether hyperglycemia-induced loss of nitric oxide in the LSEC can drive lipid accumulation in the hepatocyte. We cocultured primary mouse hepa-tocytes with LSECs under low glucose and high glucose in medium without added fatty acids.

Hyperglycemia raised hepatocyte lipid accumulation to 190% of the low glucose level (P = 0.036; n = 3) (Fig. 5a, 5b). To ask whether loss of nitric oxide alone is sufficient, we added the nitric oxide synthase inhibitor L-NAME to cocultures under low glucose. Hepatocyte lipid accumulation rose 2.2-fold (P = 0.048; n = 3). The nitric oxide donor DETA-NONOate, applied under hyperglycemia, lowered lipid accumulation to 62% of the hyperglycemic level (P = 0.027; n = 3) and returned it to 117% of the low glucose level (P = 0.53; n = 3).

**Fig. 5.**
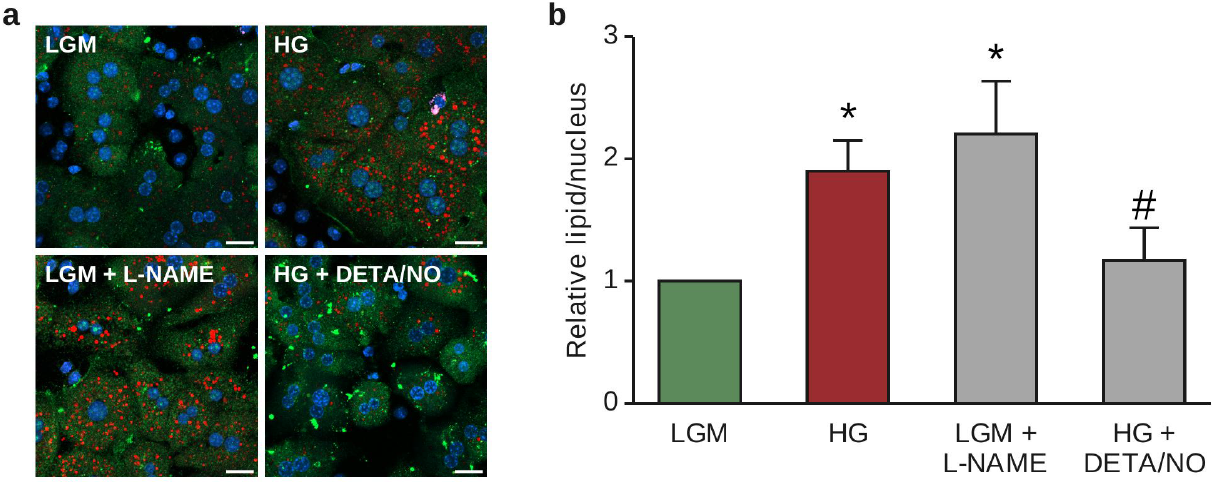
Loss of sinusoidal nitric oxide drives hepatocyte lipid accumulation. a, LSEC-hepatocyte cocultures stained for neutral lipid (LipidTOX, red), the hepatocyte marker ASGR1 (green) and DAPI (blue), in LGM, HG, LGM with the nitric oxide synthase inhibitor L-NAME, and HG with the nitric oxide donor DETA/NO. No fatty acid was added. **b,** High glucose and L-NAME increase hepatocyte lipid accumulation, and DETA/NO decreases it under high glucose. LipidTOX signal per hepatocyte nucleus, counting only nuclei within the ASGR1 mask, relative to LGM (n = 3 preparations, one-sample t-test on log2 ratios). Error bars, s.e.m. *P *<* 0.05 versus LGM. #P *<* 0.05 versus HG. Scale bars, 20 µm.

Loss of nitric oxide in the LSEC is therefore sufficient to increase hepatocyte lipid accumulation, and restoring nitric oxide under hyperglycemia reduces it.

### Lowering protein O-GlcNAcylation restores fenestrae and attenuates fibrosis in established MASH

To test whether lowering protein O-GlcNAcylation reverses the features of established MASH, mice were fed a Western diet (WD), high in fat, fructose and cholesterol, for 30 weeks, then treated with OSMI-1 or vehicle every other day for 4 weeks (Fig. 6a).

**Fig. 6.**
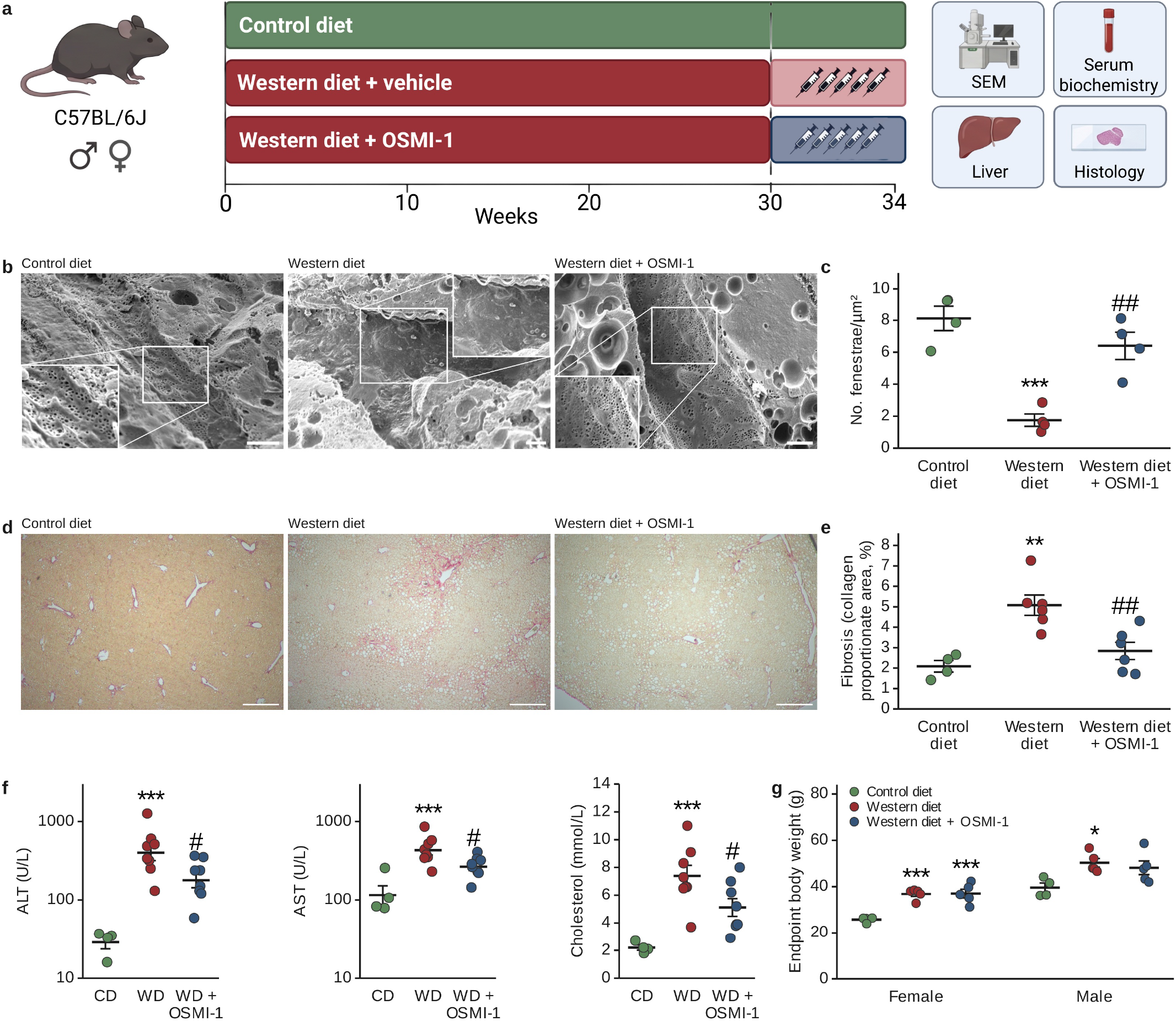
Inhibition of protein O-GlcNAcylation restores sinusoidal fenestrae and attenuates fibrosis in mice with established MASH. **a,** Study design. Control diet or Western diet (WD) for 30 weeks, then OSMI-1 or vehicle for 4 weeks, injected every other day. **b,** Representative scanning electron micrographs of sinusoidal endothelium. **c,** Number of fenestrae per square micrometer. Fenestrae are lost on the Western diet and restored by OSMI-1 (n = 4 mice per group, one-way ANOVA with Tukey’s test). **d,** Sirius red sections. **e,** Fibrosis quantification (collagen proportionate area). Collagen is increased on the Western diet and reduced by OSMI-1 (n = 4, 6 and 6 mice, one-way ANOVA with Tukey’s test). **f,** Serum ALT, AST and cholesterol. All three are increased on the Western diet and reduced by OSMI-1 (n = 4, 8 and 8 mice, one-way ANOVA with Dunnett’s test against the Western diet group, ALT and AST log-transformed). **g,** Endpoint body weight, shown separately for females and males. Body weight is increased on the Western diet and is not changed by OSMI-1 (n = 8, 10 and 10 mice, one-way ANOVA with Tukey’s test within each sex). All tests were performed on per-animal values. Each dot is one mouse. Line, mean. Error bars, s.e.m. *P *<* 0.05, **P *<* 0.01, ***P *<* 0.001 versus control diet. #P *<* 0.05, ##P *<* 0.01 versus Western diet. Scale bars, 3 µm (b) and 100 µm (d). Panel **a** created in BioRender. Levy, G. (2026) https://BioRender.com/ok5krma

Scanning electron microscopy of the sinusoidal endothelium *in vivo* showed loss of fenestrae in Western diet animals and their restoration in OSMI-1 treated animals (Fig. 6b). Fenestrae frequency fell from 8.12 per µm^2^ on control diet to 1.75 per µm^2^ on the Western diet (P = 0.00053; n = 4 animals), and OSMI-1 raised it to 6.40 per µm^2^ (P = 0.0027; n = 4), restoring fenestrae to a level not significantly different from the control diet (P = 0.24) (Fig. 6c). Fibrosis, quantified as collagen proportionate area, rose from 2.09% on control diet to 5.08% on the Western diet (P = 0.0016; n = 4 and 6), and OSMI-1 reduced it to 2.84% (P = 0.0059; n = 6), a level not significantly different from the control diet (P = 0.51) (Fig. 6d, 6e). Serum ALT, AST and cholesterol all rose on the Western diet (P = 3.5 × 10^-6^, P = 9.3 × 10^-5^ and P = 4.1 × 10^-4^) and all fell with OSMI-1 treatment (P = 0.030, P = 0.048 and P = 0.041), while remaining above the control diet level (Fig. 6f). The Western diet raised body weight in both sexes (P = 0.00048 and P = 0.025 for females and males), and OSMI-1 did not change it (P = 0.996 and P = 0.79), so its liver effects occurred independently of weight loss (Fig. 6g).

Lowering protein O-GlcNAcylation therefore restores sinusoidal fenestrae, attenuates fibrosis and lowers markers of liver injury in animals with established disease.

## Discussion

The results presented here identify the liver sinusoidal endothelial cell as a target of hyperglycemia, and define a pathway linking glucose, capillarization of the LSEC and the accumulation of lipid in the hepatocyte. The glucotoxicity lesion in the LSEC is cell-autonomous and arises through a mechanism already described in diabetic retinopathy, nephropathy and neuropathy, in which unrestricted glucose entry inhibits GAPDH and shunts the proximal intermediates into the pathways of glucotoxicity. We measured the mechanism in the LSEC, from poly-ADP-ribosylation of GAPDH to defenestration (Fig. 1–3).

Exposure of the hepatic sinusoid to hyperglycemia begins years before diabetes can be diagnosed^22–24^. Postprandial glucose reaches the LSEC at portal concentrations rather than systemic, and in insulin resistance those excursions are larger^9,10^. After a meal, portal glucose exceeds arterial glucose, creating a negative arterioportal gradient^9^. That gradient exposes the LSEC to glucose concentrations higher than the systemic circulation reaches. The fasting glucose and HbA1c thresholds for diagnosing diabetes were set at the systemic levels at which retinopathy becomes detectable^56,57^. The retina, along with the kidney, is perfused by systemic blood. Our results place the mechanism that produces retinopathy and nephropathy in the hepatic sinusoid. The diagnostic thresholds do not measure the portal circulation, where glucose concentrations are highest.

Metabolic liver disease is understood to begin in the hepatocyte, where lipid delivery exceeds its lipid-handling capacity and the resulting lipotoxicity drives inflammation and fibrosis^58–60^. Damage-associated molecular patterns and cytokines released by the injured hepatocyte then activate the non-parenchymal cells of the liver^61,62^. Hepatocyte-derived extracellular vesicles have been reported to deliver OGT to the LSEC in MASLD^63^. We show that hyperglycemia capillarizes the LSEC in isolation (Fig. 1), so the endothelial lesion does not require parenchymal injury. Inhibition of nitric oxide synthase in the LSEC raised hepatocyte lipid accumulation in coculture (Fig. 5), so the endothelium contributes to steatosis. The hepatocyte stores glucose as glycogen and converts it to lipid, which it exports in lipoproteins^64^. The LSEC has none of these routes, cannot limit glucose entry, and is the first cell exposed to portal blood.

Of the four pathways of glucotoxicity, only inhibition of hexosamine flux with DON significantly restored porosity under hyperglycemia (Fig. 2). Raising protein O-GlcNAcylation under low glucose was itself sufficient to defenestrate the LSEC (Fig. 2f). Inhibition of aldose reductase with zopolrestat, advanced glycation with pyridoxamine, and protein kinase C β with ruboxistaurin increased porosity without reaching significance. In the classical diabetic complications these pathways are activated by a single upstream event^6,7^. Aldose reductase consumes NADPH, so inhibiting it preserves glutathione regeneration^7^. Advanced glycation end products quench nitric oxide^65^, and protein kinase C phosphorylates eNOS^66^ at the inhibitory site Thr^495^. A contribution from any of them cannot be excluded. Hexosamine flux to protein O-GlcNAcylation, however, was sufficient to capillarize the LSEC.

In the LSEC the hexosamine biosynthetic pathway has a second input, one that supplies GlcNAc independently of glucose uptake. GlcNAc released by lysosomal degradation of cleared hyaluronan is diverted toward protein O-GlcNAcylation when glycolytic disposal is restricted (Fig. 4). Indeed, serum hyaluronan rises with fibrosis stage and is measured in clinical fibrosis tests^67–69^. Since hyaluronan is cleared by the LSEC, its accumulation in serum is a predicted consequence of capillarization. Under hyperglycemia the LSEC’s clearance function therefore supplies substrate to the pathway that capillarizes it.

Postprandial glucose and insulin resistance both rise with age, independently of adiposity^70,71^. Pseudocapillarization, the loss of fenestrae that accompanies aging^72,73^, has no established cause. A glucose-driven loss of nitric oxide is one candidate mechanism. Several agents that restore fenestrations in aged animals act at steps of this cascade^40,74–76^. If pseudocapillarization shares this mechanism, lowering protein O-GlcNAcylation should restore fenestrae in aged animals.

Inhibition of protein O-GlcNAcylation in mice with established MASH restored LSEC fenestrae and attenuated fibrosis (Fig. 6). OSMI-1 also lowered protein O-GlcNAcylation in the human LSEC (Supplementary Fig. 3). This identifies the hexosamine biosynthetic pathway and protein O-GlcNAcylation as potential therapeutic targets in MASLD.

Diabetes predicts progression to cirrhosis and hepatocellular carcinoma, and glycemic control reduces fibrosis independently of body weight^2–5^. Lean patients also develop steatotic liver disease, and higher fasting glucose is independently associated with the lean phenotype^77,78^. These observations point to hyperglycemia, yet the injury it produces in the liver is unknown^1^. A glucose-driven lesion of the sinusoidal endothelium would account for them. It would also explain why glycemic control improves fibrosis independently of body weight, just as it protects the retina and the kidney^79,80^.

We propose that the hepatic sinusoid is a site of microvascular complications of hyperglycemia, alongside the retina, the kidney and the peripheral nerve. Capillarization of the LSEC is a mechanism by which glycemia promotes steatotic liver disease, and the basis for a therapeutic strategy directed at the endothelium.

## Materials and Methods

### Animals and ethics

Animal protocols were approved by the institutional authority at UiT The Arctic University of Norway and licensed by the Norwegian Food Safety Authority (Mattilsynet; FOTS ID 31882; *in vitro* approval 06/26), in compliance with EU Directive 2010/63/EU and reported following the ARRIVE guidelines. Primary cell isolations were performed using male BALB/c mice. *In vivo* studies were performed using male and female C57BL/6JRj mice (Janvier Labs), enrolled at 8 weeks of age (18 to 22 g). Mice were housed in Eurostandard type IIIH cages with environmental enrichment and provided ad libitum access to food and water. Mice were allocated to control diet (Research Diets D09100304; 10% kcal fat, 0% fructose, 0% cholesterol) (n = 8; 4 female, 4 male) or a Western diet (Research Diets D09100301; 40% fat, mostly Primex, 20% fructose, 2% cholesterol) (n = 20; 10 female, 10 male). After 30 weeks on diet, Western diet mice were randomly allocated by blinded selection to OSMI-1 (n = 10; 5 female, 5 male) or vehicle (n = 10; 5 female, 5 male) and received intraperitoneal injections of OSMI-1 (10 mg/kg, formulated in 4.5% DMSO, 5% Tween 80 and 90.5% saline) or vehicle every second day for 4 weeks^81^. At the end of the study, body weight was recorded and mice were euthanized for liver tissue and serum collection.

### Isolation of primary mouse liver sinusoidal endothelial cells and hepatocytes

LSECs and hepatocytes were isolated from the same liver^82^. Liver was perfused via the portal vein with calcium free HEPES buffered saline (142 mM NaCl,6.7 mM KCl, 9.6 mM HEPES, pH adjusted with NaOH), then the same buffer with 4.76 mM CaCl_2_ and 0.02 mg/mL Liberase TM (Merck, 5401127001) until parenchyma dissociated. Hepatocytes were recovered by differential centrifugation, purified on 45% Percoll gradient (Sigma Aldrich, GE17-0891-01), yielding above 80% viability, and seeded on fibronectin-coated surfaces in Williams’ medium E (Thermo Fisher, A1217601) supplemented with dexamethasone (0.1 mM), L-glutamine (2 mM), insulin (86 nM), penicillin/streptomycin (50 U/mL, 50 µg/mL), and foetal bovine serum (FBS) (10%). LSECs were purified from the nonparenchymal fraction by magnetic activated cell sorting using mouse anti-CD146 (LSEC) MicroBeads (Miltenyi Biotec, 130-092-007) on LS Columns (Miltenyi Biotec, 130-042-401) with a QuadroMACS Separator, and seeded on fibronectin in low glucose (5.5 mM) DMEM (Merck, D5546) supplemented with L-glutamine (2 mM), penicillin/ streptomycin (50 U/mL, 50 µg/mL), and 2 mg/mL human serum albumin (Alburex 20, 200 g/L, CSL Behring). Cultures were washed 30 to 40 min after seeding, removing debris and non-adherent cells. This preparation yields 95-98% purity, assessed as fenestrated cells by scanning electron microscopy^82^. All experiments were completed within 24 h of seeding.

### Mouse and human LSEC treatments

Purified mouse LSECs were incubated 24 h at 37°C, 5% CO_2_, 5% O_2_ in low glucose (5.5 mM) or high glucose (30 mM), with or without the following compounds: D-glucosamine hydrochloride 5 mM (G4875, Merck), Thiamet-G 5 µM (SML0244, Merck); OSMI-1 10 µM (SML1621, Merck); 6-diazo-5-oxo-L-norleucine (DON) 20 µM (D2141, Merck); heptelidic acid 10 µM (ab144269, Abcam); zopolrestat 10 µM (Z4527, Merck); ruboxistaurin 400 nM (TA9947273119, Merck); pyridoxamine dihydrochloride 1 mM (P9158, Merck); Y-27632 10 µM (Y0503, Merck); DETA-NONOate 50 µM (82120, BioNordika); L-NAME 100 µM (483125, Merck); ODQ 10 µM (HY-101255, Nordic Biosite); 8-bromo-cGMP 100 µM (105-B-2174, Nordic Biosite); hyaluronan (medium MW, GLR004, R&D Systems) 400 µg/mL at start plus 200 µg/mL at 20 h. OSMI-1, DON and zopolrestat added 30 min before high glucose, and ODQ 30 min before DETA-NONOate. D-mannitol (M4125, Merck) 24.5 mM or 5 mM controlled osmolarity for the high glucose and glucosamine arms respectively. Solvent concentrations were matched across treatments, with DMSO added to control wells at the concentration present in the corresponding treatment.

Primary human LSECs (5000, ScienCell) were seeded onto fibronectin-coated 6-well plates and cultured in EBM-2 Endothelial Cell Growth Basal Medium (CC-3156, Lonza; baseline 5.5 mM glucose) supplemented with endothelial growth factors, penicillin/streptomycin, and 5% fetal bovine serum (FBS). Upon reaching 80% confluence, cells were switched to serum-free medium and incubated for 24 h at 37°C, 5% CO_2_, and 20% O_2_ under treatment with or without D-glucosamine hydrochloride (5 mM) and Thiamet-G (5 µM), in the presence or absence of OSMI-1 (10 µM).

### LSEC-hepatocyte co-culture and hepatocyte lipid quantification

Hepatocytes and LSECs from the same liver were co-cultured at 1:1 ratio in 1:1 Williams’ medium E/DMEM, hepatocytes seeded first then LSECs, in serum and fatty acid free medium. Co-cultures were incubated 24 h at 37°C, 5% CO_2_, 20% O_2_ in low glucose (5.5 mM), high glucose (30 mM), low glucose plus 100 µM L-NAME, or high glucose plus 50 µM DETA-NONOate. Low glucose with and without 24.5 mM D-mannitol gave indistinguishable results, thus later treatments used mannitol matched low glucose. Co-cultures were fixed in 4% paraformaldehyde/PBS for 10 min and blocked for 60 min in 3% BSA/PBS without detergent. Hepatocytes were identified with rabbit anti-ASGR1 (Thermo Fisher, PA5-32030) 4 µg/mL, 1% BSA/PBS, 1 h, room temperature, then donkey anti-rabbit Alexa Fluor 488 (Thermo Fisher, A21206) same concentration). Neutral lipid was detected using HCS LipidTOX Red (Invitrogen, H34476), and nuclei using DAPI. In Fiji, LipidTOX signal within the ASGR1 mask was summed and divided by the number of nuclei within that mask.

### Scanning electron microscopy

#### Cultured LSECs

LSECs in 16-well CultureWell coverglass (Grace Bio-Labs/Merck; GBL112358) were fixed in 1% glutaraldehyde, 4% formaldehyde, PHEM buffer, followed by 40 min incubation in 1% tannic acid/PHEM, post fixed for maximum 30 min in 1% osmium tetroxide/ddH_2_O, dehydrated in 30%, 60%, 90% ethanol (5 min each), four 5 min absolute ethanol changes, then dried two 10 min in hexamethyldisilazane (HMDS) (Merck, 440191) exchanges.

#### Liver tissue

Liver tissue was prepared as described^83^. Post euthanasia, animals were perfusion fixed via left cardiac ventricle with PBS (pH 7.4) then 4% formaldehyde in PBS (pH 7.4). Tissue blocks (1-2 mm^3^) from fixed left lobe edges were immersed in 1% glutaraldehyde/4% formaldehyde in PHEM buffer, processed as cultured cells to absolute ethanol, dropped into liquid nitrogen, cracked with razor and hammer to expose sinusoid luminal surfaces, returned to ethanol, and dried in HMDS.

Dried specimens were mounted on aluminum stubs with silver glue and sputter coated with 10 nm gold/palladium alloy. Specimens were imaged on a Zeiss Gemini 300 scanning electron microscope. Fifteen to twenty cells were imaged per preparation, from five or more randomly selected regions of the well, and 15 to 20 fields per animal for tissue.

### Quantification of fenestrations

Fenestrations were segmented with a StarDist model, which represents objects as star-convex polygons, trained on SEM images of LSEC^84,85^. From the segmentation we derived, for each cell, the number of fenestrae per µm^2^ of cell area, and porosity, the percentage of the cell area they occupy^86^. Segmentation used the StarDist plugin in Fiji with the previously trained LSEC fenestrae model^84^. Cell area was outlined manually in Fiji.

### Quantitative reverse-transcriptase PCR

LSEC isolated from 5 mice were established in fibronectin-coated 48 well plates (Sarstedt). Cellular RNA was isolated and reverse transcribed using the TaqMan Fast Advanced Cells to CT Kit (Thermo Fisher, A35374). cDNA was pre amplified with TaqMan PreAmp Master Mix (Thermo Fisher, 4391128). qPCR used TaqMan Gene Expression Assays on a QuantStudio 5 Real Time PCR System (Applied Biosystems). 10 µL reactions contained 2.5 µL cDNA, 0.5 µL TaqMan Gene Expression Assay, 5 µL TaqMan Fast Advanced Master Mix (Thermo Fisher, 4444556). Cycling: UDG activation 50°C/2 min, denaturation 95°C/20 s; 40 cycles of 95°C/1 s, 60°C/20 s. Assay identifiers are listed in Supplementary Table 2. Quantification cycles were averaged across the technical replicates of each assay in each sample. Relative mRNA expression was normalized to the geometric mean of four reference genes, *Rpl13a, Ppia, B2m* and *Hprt*, measured in the same sample. Transcripts in high glucose are expressed relative to low glucose in the same animal as 2^-DDCq. Glucose transporter transcripts are compared with one another as a percentage of the geometric mean of the reference genes, 100 × 2^-DCq, pooled across samples.

### Western blotting

Cells cultured in 6-well plates were washed with ice-cold PBS containing 10 µM Thiamet-G and lysed in modified RIPA buffer supplemented with 10 µM Thiamet-G, protease and Halt phosphatase inhibitor cocktails, 1 mM sodium orthovanadate, 10 mM N-ethylmaleimide, and 10 µg/mL pepstatin A. Lysates were cleared by centrifugation (10,000 ×g, 15 min, 4°C), and protein concentration was determined using the Micro BCA Protein Assay Kit (Thermo Fisher, 23235).

Denatured protein samples (2.5 µg per lane) were separated on NuPAGE 4–12% Bis-Tris gradient gels in 1x MOPS buffer and transferred to 0.45 µm PVDF membranes using a semi-dry blotting system. Membranes were blocked in milk-free EveryBlot Blocking Buffer and probed for global O-GlcNAcylation overnight at 4°C with RL2 (Thermo Fisher, 2 µg/mL), followed by goat antimouse IgG-HRP. Chemiluminescent detection was performed using SuperSignal West Pico PLUS substrate and imaged on an ImageQuant LAS4000. Membranes were subsequently stripped and reprobed for β-actin (Abcam, ab8227) as a loading reference.

Global O-GlcNAc levels were quantified by integrating signal over the full molecular weight range of each lane and dividing by the β-actin band of the same lane. Lanes were then expressed relative to the mannitol matched low glucose lane of the same animal on the same membrane.

### Proximity ligation assay

In situ proximity ligation (UnFold probe^87^) was performed on LSECs cultured in 16 well CultureWell removable chambered coverglass, using NaveniFlex Cell MR Red kit (Navinci Diagnostics, 60025) per manufacturer’s instructions with an additional blocking step using ReadyProbes Mouse-on-Mouse IgG Blocking Solution (Thermo Fisher, R37621) prior to the Naveni Block. Antibody pairs used were rabbit anti-eNOS, 1.5 µg/mL (Abcam, ab199956, EPR19296) with mouse anti-O-GlcNAc RL2, 2 µg/mL (Thermo Fisher, MA1-072), and rabbit anti-GAPDH, 0.1 µg/mL (Thermo Fisher, PA1-16777) with mouse anti-PAR 10H, 2 µg/mL (Enzo/AH Diagnostics, ALX-804-220-R100). Proximity ligation signal was segmented in the red channel in Fiji at a fixed threshold applied to every image, and is reported as signal area per nucleus^88^, averaged across fields within an animal.

### GAPDH activity assay

GAPDH activity was measured with a colorimetric kit (Abcam, ab204732). Cell cultures were washed in ice-cold PBS, resuspended in 100 µL ice-cold Assay Buffer 4 per well in 3 wells of a 6-well plate, pooled, iced for 10 min, and cleared at 10,000 × g for 5 min, 4°C. Supernatants were assayed in triplicate against NADH standard curve in a 96 well plate. Absorbance at 450 nm was read kinetically and activity was taken as the slope over the linear phase, corrected for the slope of enzyme-free control wells, converted using the NADH standard curve of the same plate, and expressed as nmol NADH/min per million cells.

### Nitric oxide measurement

Nitric oxide was measured with DAF-FM diacetate (Thermo Fisher, D23844)^89^ in LSECs at 70 to 80% confluence, treated for 24 h. In the final hour, the cells were washed twice in HBSS (Ca/Mg), incubated with 5 µM DAF-FM diacetate including the treatments at 37°C, then the buffer replaced with HBSS containing 2 µM Hoechst 33342 for 20 min and imaged in HBSS. DAF-FM signal was summed over each field above a local background threshold and divided by the number of nuclei in that field, obtained by Otsu thresholding of the nuclear channel in Fiji. Each preparation is expressed relative to its own low glucose mean.

### Immunofluorescence

For F-actin staining, the cells were fixed in 4% PFA (10 min), 0.1% Triton X-100/PBS (10 min), 3% BSA/PBS block, phalloidin-Alexa Fluor 488 (Thermo Fisher, A12379) 1:40 in 1% BSA/PBS for 30 min, and DAPI. F-actin is reported as mean phalloidin intensity per nucleus, measured in Fiji.

For global O-GlcNAc, cells were fixed, permeabilized and blocked sequentially with 5% normal donkey serum (30 min) then Mouse-on-Mouse blocking reagent for 1 h. RL2 antibody (4 µg/ml) was applied overnight at 4°C followed by 1 h incubation at room temperature with donkey anti mouse DyLight 488 (Thermo Fisher, SA5-10166), same concentration.

Fixed-cell immunofluorescence images were automatically acquired on a Zeiss LSM 800 confocal laser scanning microscope using automated grid tiling. For each preparation, 20 tiled regions were imaged per well. Acquisition settings were held constant across all preparations within an experiment.

### Untargeted metabolomics

For metabolomics, LSECs were harvested as dry pellets. Spent medium was aspirated, cells washed three times in cold PBS (for less than 30 s), and cells scraped and collected into 1 mL PBS, centrifuged at 1,000 × g, for 3 min, 4°C. The supernatant removed completely and pellets snap frozen in liquid nitrogen, stored at -80°C and shipped on dry ice to Metabolon, Inc. (Morrisville, NC, USA) for UHPLC-MS/MS on the Global Discovery platform. The metabolites were identified against authenticated standards by retention index, accurate mass and MS/MS fragmentation. Peaks were quantified using area under the curve. The abundance of each biochemical was normalized to the cellular DNA concentration of the corresponding sample, missing values were imputed with the minimum observed value for each compound, values were log transformed, and one-way ANOVA was used to identify biochemicals that differed between experimental groups.

### Histochemistry

Paraffin sections of 4 µm thickness were cut from perfused fixed livers, deparaffinized in xylene, rehydrated through a graded ethanol series, and stained with Picro Sirius red (Abcam, ab150881). Four to five fields per animal were imaged in brightfield at 4x (1.61 µm/pixel) on a Nikon inverted microscope with a Zeiss AxioCam MRc camera. Collagen was quantified in Fiji^90^ by HSB color thresholding, excluding the liver capsule. Fibrosis was quantified as collagen proportionate area, the thresholded area as a percentage of tissue area, averaged per mouse.

### Serum biochemistry

Serum alanine aminotransferase, aspartate aminotransferase and cholesterol were measured using Atellica analyzer at Sentrallaboratoriet, NMBU Veterinaer-hogskolen.

### Statistics

The unit of analysis was the cell preparation for experiments *in vitro* and the animal for experiments *in vivo*. Fields, wells and tissue pieces were averaged within that unit before comparison.

Ratios formed within an animal were tested on log2-transformed values by one-sample t test against no change. Absolute measures in two arms were compared by two-tailed unpaired t test on preparation means. Comparisons across more than two arms used one-way ANOVA followed by Dunnett’s test against the stated control arm.

*In vivo* endpoints were compared by one-way ANOVA on per-animal values, with fields or images averaged within each animal first, followed by Tukey’s test. Serum measures were compared by Dunnett’s test against the Western diet group, since only comparisons with that group are reported for the serum measures. ALT and AST were log-transformed before testing, and body weight was analyzed within each sex.

Metabolomics data were analyzed by Metabolon as described above and were not reanalyzed here. Exact two-sided P values are reported throughout and P *<* 0.05 was considered significant.

## Acknowledgments

The authors express special thanks to Karolina Szafranska and Jaione Simón-Santamaría for their valuable assistance with whole-body perfusion and scanning electron microscopy preparations. We extend our sincere gratitude to the staff at the Section for Comparative Medicine (Amalie H. Andersen, Carina Sørensen, Hallvard Haugen, Izabela J. Szkotak, Helene S.W. Jeppesen, Remi A.H. Osnes, Lorenzo Ragazzi) for their essential support with animal protocol licensing and intraperitoneal injections. We also acknowledge the members of the Advanced Microscopy Platform (Tom-Ivar Eilertsen, Kenneth B. Laesen, Randi Olsen, Aud-Malin K. Hovd, Roy A. Lyså) for their expert technical assistance, facilities, and guidance throughout this study.

## Author contributions

G.L. conceived the study, designed the experiments, analyzed the data, and wrote the manuscript. C.I.Ø. secured regulatory approval (FOTS license), contributed to experimental design, performed cell isolation and imaging experiments, and participated in data analysis. G.D. managed chemical and reagent preparation, performed Western blotting, tissue sectioning, and staining, and assisted with MASH mouse liver sampling, cell isolation, and experimental logistics. R.L. designed and performed the qRT-PCR assays, executed the majority of liver perfusions, assisted with MASH mouse tissue sampling, and performed the GAPDH activity assays. All authors reviewed and approved the final manuscript.

## References

1 Cusi, K. et al. Metabolic Dysfunction-Associated Steatotic Liver Disease (MASLD) in People With Diabetes: The Need for Screening and Early Intervention. A Consensus Report of the American Diabetes Association. Diabetes Care 48, 1057–1082 (2025).

2 Alexander, M. et al. Risks and clinical predictors of cirrhosis and hepatocellular carcinoma diagnoses in adults with diagnosed NAFLD: real-world study of 18 million patients in four European cohorts. BMC Med 17, 95 (2019).

3 Kanwal, F. et al. Effect of Metabolic Traits on the Risk of Cirrhosis and Hepatocellular Cancer in Nonalcoholic Fatty Liver Disease. Hepatology 71, 808–819 (2020).

4 Hamaguchi, E. et al. Histological course of nonalcoholic fatty liver disease in Japanese patients: tight glycemic control, rather than weight reduction, ameliorates liver fibrosis. Diabetes Care 33, 284–6 (2010).

5 Sako, S. et al. Trajectories of Liver Fibrosis and Gene Expression Profiles in Nonalcoholic Fatty Liver Disease Associated With Diabetes. Diabetes 72, 1297–306 (2023).

6 Brownlee, M. Biochemistry and molecular cell biology of diabetic complications. Nature 414, 813–20 (2001).

7 Brownlee, M. The pathobiology of diabetic complications: a unifying mechanism. Diabetes 54, 1615–25 (2005).

8 Nishikawa, T., Edelstein, D. & Brownlee, M. The missing link: a single unifying mechanism for diabetic complications. Kidney Int Suppl 77, S26–30 (2000).

9 Björkman, O., Eriksson, L. S., Nyberg, B. & Wahren, J. Gut exchange of glucose and lactate in basal state and after oral glucose ingestion in postoperative patients. Diabetes 39, 747–51 (1990).

10 Lund, B., Schmidt, A. & Deckert, T. Portal and cubital serum insulin during oral, portal and cubital glucose tolerance tests. Acta Med Scand 197, 275–81 (1975).

11 Felig, P., Wahren, J. & Hendler, R. Influence of oral glucose ingestion on splanchnic glucose and gluconeogenic substrate metabolism in man. Diabetes 24, 468–75 (1975).

12 Yosefy, C. Hyperglycaemia and its relation to cardiovascular morbidity and mortality: has it been resolved? Acta Diabetol 40 Suppl 2, S380–8 (2003).

13 DeFronzo, R. A., Bonadonna, R. C. & Ferrannini, E. Pathogenesis of NIDDM. A balanced overview. Diabetes Care 15, 318–68 (1992).

14 Sørensen, K. K., Simon-Santamaria, J., McCuskey, R. S. & Smedsrød, B. Liver Sinusoidal Endothelial Cells. Compr Physiol 5, 1751–74 (2015).

15 DeLeve, L. D. & Maretti-Mira, A. C. Liver Sinusoidal Endothelial Cell: An Update. Semin Liver Dis 37, 377–387 (2017).

16 Seternes, T., Sørensen, K. & Smedsrød, B. Scavenger endothelial cells of vertebrates: a nonperipheral leukocyte system for high-capacity elimination of waste macromolecules. Proc Natl Acad Sci U S A 99, 7594–7 (2002).

17 Sørensen, K. K. et al. The scavenger endothelial cell: a new player in homeostasis and immunity. Am J Physiol Regul Integr Comp Physiol 303, R1217–30 (2012).

18 Bhandari, S., Larsen, A. K., McCourt, P., Smedsrød, B. & Sørensen, K. K. The Scavenger Function of Liver Sinusoidal Endothelial Cells in Health and Disease. Front Physiol 12, 757469 (2021).

19 Hammoutene, A. & Rautou, P. E. Role of liver sinusoidal endothelial cells in non-alcoholic fatty liver disease. J Hepatol 70, 1278–1291 (2019).

20 DeLeve, L. D. Liver sinusoidal endothelial cells in hepatic fibrosis. Hepatology 61, 1740–6 (2015).

21 Natarajan, V., Harris, E. N. & Kidambi, S. SECs (Sinusoidal Endothelial Cells), Liver Microenvironment, and Fibrosis. Biomed Res Int 2017, 4097205 (2017).

22 Hanefeld, M., Sulk, S., Helbig, M., Thomas, A. & Köhler, C. Differences in Glycemic Variability Between Normoglycemic and Prediabetic Subjects. J Diabetes Sci Technol 8, 286–290 (2014).

23 Ding, C. et al. Regulation of glucose metabolism in nondiabetic, metabolically obese normal-weight Asians. Am J Physiol Endocrinol Metab 314, E494–E502 (2018).

24 Kashiwagi, K. et al. Assessment of glycemic variability and lifestyle behaviors in healthy nondiabetic individuals according to the categories of body mass index. PLoS One 18, e0291923 (2023).

25 Kaiser, N. et al. Differential regulation of glucose transport and transporters by glucose in vascular endothelial and smooth muscle cells. Diabetes 42, 80–9 (1993).

26 Marshall, S., Bacote, V. & Traxinger, R. R. Discovery of a metabolic pathway mediating glucose-induced desensitization of the glucose transport system. Role of hexosamine biosynthesis in the induction of insulin resistance. J Biol Chem 266, 4706–12 (1991).

27 Peterson, S. B. & Hart, G. W. New insights: A role for O-GlcNAcylation in diabetic complications. Crit Rev Biochem Mol Biol 51, 150–61 (2016).

28 Torres, C. R. & Hart, G. W. Topography and polypeptide distribution of terminal N-acetylglucosamine residues on the surfaces of intact lymphocytes. Evidence for O-linked GlcNAc. J Biol Chem 259, 3308–17 (1984).

29 Kreppel, L. K., Blomberg, M. A. & Hart, G. W. Dynamic glycosylation of nuclear and cytosolic proteins. Cloning and characterization of a unique O-GlcNAc transferase with multiple tetratricopeptide repeats. J Biol Chem 272, 9308–15 (1997).

30 Ortiz-Meoz, R. F. et al. A small molecule that inhibits OGT activity in cells. ACS Chem Biol 10, 1392–7 (2015).

31 Dong, D. L. & Hart, G. W. Purification and characterization of an O-GlcNAc selective N-acetyl-beta-D-glucosaminidase from rat spleen cytosol. J Biol Chem 269, 19321–30 (1994).

32 Gao, Y., Wells, L., Comer, F. I., Parker, G. J. & Hart, G. W. Dynamic O-glycosylation of nuclear and cytosolic proteins: cloning and characterization of a neutral, cytosolic beta-N-acetylglucosaminidase from human brain. J Biol Chem 276, 9838–45 (2001).

33 Yuzwa, S. A. et al. A potent mechanism-inspired O-GlcNAcase inhibitor that blocks phosphorylation of tau in vivo. Nat Chem Biol 4, 483–90 (2008).

34 Francis, S. H., Busch, J. L., Corbin, J. D. & Sibley, D. cGMP-dependent protein kinases and cGMP phosphodiesterases in nitric oxide and cGMP action. Pharmacol Rev 62, 525–63 (2010).

35 Smolenski, A., Poller, W., Walter, U. & Lohmann, S. M. Regulation of human endothelial cell focal adhesion sites and migration by cGMP-dependent protein kinase I. J Biol Chem 275, 25723–32 (2000).

36 Sawada, N. et al. cGMP-dependent protein kinase phosphorylates and inactivates RhoA. Biochem Biophys Res Commun 280, 798–805 (2001).

37 Zapotoczny, B., Szafranska, K., Lekka, M., Ahluwalia, B. S. & McCourt, P. Tuning of Liver Sieve: The Interplay between Actin and Myosin Regulatory Light Chain Regulates Fenestration Size and Number in Murine Liver Sinusoidal Endothelial Cells. Int J Mol Sci 23, 9850 (2022).

38 Venkatraman, L. & Tucker-Kellogg, L. The CD47-binding peptide of thrombospondin-1 induces defenestration of liver sinusoidal endothelial cells. Liver Int 33, 1386–97 (2013).

39 Di Martino, J. et al. Actin Depolymerization in Dedifferentiated Liver Sinusoidal Endothelial Cells Promotes Fenestrae Re-Formation. Hepatol Commun 3, 213–219 (2019).

40 Hunt, N. J. et al. Manipulating fenestrations in young and old liver sinusoidal endothelial cells. Am J Physiol Gastrointest Liver Physiol 316, G144–G154 (2019).

41 Aulak, K. S. et al. Specific O-GlcNAc modification at Ser-615 modulates eNOS function. Redox Biol 36, 101625 (2020).

42 Masaki, N. et al. O-GlcNAcylation Mediates Glucose-Induced Alterations in Endothelial Cell Phenotype in Human Diabetes Mellitus. J Am Heart Assoc 9, e014046 (2020).

43 Du, X. L. et al. Hyperglycemia inhibits endothelial nitric oxide synthase activity by posttranslational modification at the Akt site. J Clin Invest 108, 1341–8 (2001).

44 Musicki, B., Kramer, M. F., Becker, R. E. & Burnett, A. L. Inactivation of phosphorylated endothelial nitric oxide synthase (Ser-1177) by O-GlcNAc in diabetes-associated erectile dysfunction. Proc Natl Acad Sci U S A 102, 11870–5 (2005).

45 Amano, M., Nakayama, M. & Kaibuchi, K. Rho-kinase/ROCK: A key regulator of the cytoskeleton and cell polarity. Cytoskeleton (Hoboken) 67, 545–54 (2010).

46 Smedsrød, B., Pertoft, H., Eriksson, S., Fraser, J. R. & Laurent, T. C. Studies in vitro on the uptake and degradation of sodium hyaluronate in rat liver endothelial cells. Biochem J 223, 617–26 (1984).

47 McCourt, P. A., Smedsrød, B. H., Melkko, J. & Johansson, S. Characterization of a hyaluronan receptor on rat sinusoidal liver endothelial cells and its functional relationship to scavenger receptors. Hepatology 30, 1276–86 (1999).

48 Campbell, P. et al. N-acetylglucosamine-6-phosphate deacetylase in hepatocytes, Kupffer cells and sinusoidal endothelial cells from rat liver. Hepatology 11, 199–204 (1990).

49 Kroef, V. et al. GFPT2/GFAT2 and AMDHD2 act in tandem to control the hexosamine pathway. Elife 11, e69223 (2022).

50 Kim, P. K. et al. Hyaluronic acid fuels pancreatic cancer cell growth. Elife 10, e62645 (2021).

51 Arreola, R., Valderrama, B., Morante, M. L. & Horjales, E. Two mammalian glucosamine-6-phosphate deaminases: a structural and genetic study. FEBS Lett 551, 63–70 (2003).

52 Cordero-Herrera, I. et al. AMP-activated protein kinase activation and NADPH oxidase inhibition by inorganic nitrate and nitrite prevent liver steatosis. Proc Natl Acad Sci U S A 116, 217–226 (2019).

53 Fullerton, M. D. et al. Single phosphorylation sites in Acc1 and Acc2 regulate lipid homeostasis and the insulin-sensitizing effects of metformin. Nat Med 19, 1649–54 (2013).

54 García-Villafranca, J., Guillén, A. & Castro, J. Involvement of nitric oxide/cyclic GMP signaling pathway in the regulation of fatty acid metabolism in rat hepatocytes. Biochem Pharmacol 65, 807–12 (2003).

55 Li, Y. et al. AMPK phosphorylates and inhibits SREBP activity to attenuate hepatic steatosis and atherosclerosis in diet-induced insulin-resistant mice. Cell Metab 13, 376–388 (2011).

56 Colagiuri, S. et al. Glycemic thresholds for diabetes-specific retinopathy: implications for diagnostic criteria for diabetes. Diabetes Care 34, 145–50 (2011).

57 Kumar, R., Nandhini, L. P., Kamalanathan, S., Sahoo, J. & Vivekanadan, M. Evidence for current diagnostic criteria of diabetes mellitus. World J Diabetes 7, 396–405 (2016).

58 Neuschwander-Tetri, B. A. Non-alcoholic fatty liver disease. BMC Med 15, 45 (2017).

59 Buzzetti, E., Pinzani, M. & Tsochatzis, E. A. The multiple-hit pathogenesis of non-alcoholic fatty liver disease (NAFLD). Metabolism 65, 1038–48 (2016).

60 Friedman, S. L., Neuschwander-Tetri, B. A., Rinella, M. & Sanyal, A. J. Mechanisms of NAFLD development and therapeutic strategies. Nat Med 24, 908–922 (2018).

61 Myint, M. et al. Inflammatory signaling in NASH driven by hepatocyte mitochondrial dysfunctions. J Transl Med 21, 757 (2023).

62 Arrese, M., Cabrera, D., Kalergis, A. M. & Feldstein, A. E. Innate Immunity and Inflammation in NAFLD/NASH. Dig Dis Sci 61, 1294–303 (2016).

63 Wang, Y. et al. OGT-enriched Hepatocyte-derived Extracellular Vesicles Promote Capillarization of Liver Sinusoidal Endothelial Cells in Metabolic Dysfunction-associated Steatotic Liver Disease. Cell Mol Gastroenterol Hepatol 20, 101721 (2026).

64 Rui, L. Energy metabolism in the liver. Compr Physiol 4, 177–197 (2014).

65 Bucala, R., Tracey, K. J. & Cerami, A. Advanced glycosylation products quench nitric oxide and mediate defective endothelium-dependent vasodilatation in experimental diabetes. J Clin Invest 87, 432–8 (1991).

66 Fleming, I., Fisslthaler, B., Dimmeler, S., Kemp, B. E. & Busse, R. Phosphorylation of Thr(495) regulates Ca(2+)/calmodulin-dependent endothelial nitric oxide synthase activity. Circ Res 88, E68–75 (2001).

67 Rosenberg, W. M. et al. Serum markers detect the presence of liver fibrosis: a cohort study. Gastroenterology 127, 1704–13 (2004).

68 Loomba, R. & Adams, L. A. Advances in non-invasive assessment of hepatic fibrosis. Gut 69, 1343–1352 (2020).

69 Pecoraro, V., Nascimbeni, F., Gabrielli, F., Fasano, T. & Trenti, T. Direct serum biomarkers of liver fibrosis in metabolic dysfunction-associated steatotic liver disease: A scoping review. Clin Biochem 144, 111138 (2026).

70 Shimokata, H. et al. Age as independent determinant of glucose tolerance. Diabetes 40, 44–51 (1991).

71 Qiao, Q. et al. Are insulin resistance, impaired fasting glucose and impaired glucose tolerance all equally strongly related to age? Diabet Med 22, 1476–81 (2005).

72 Le Couteur, D. G., Fraser, R., Cogger, V. C. & McLean, A. J. Hepatic pseudocapillarisation and atherosclerosis in ageing. Lancet 359, 1612–5 (2002).

73 McLean, A. J. et al. Age-related pseudocapillarization of the human liver. J Pathol 200, 112–7 (2003).

74 Hunt, N. J. et al. The Effects of Metformin on Age-Related Changes in the Liver Sinusoidal Endothelial Cell. J Gerontol A Biol Sci Med Sci 75, 278–285 (2020).

75 de Leve, S. J., Wang, X., Alavifard, H., Dara, L. & DeLeve, L. D. Reversing liver sinusoidal endothelial aging restores clearance of amyloid-beta and LDL. J Biol Chem, 113608 (2026).

76 Voloshyna, T. et al. PACA nanoparticles target and deliver sildenafil to rejuvenate aged mouse liver sinusoidal endothelial cells. Nanotheranostics 9, 155–170 (2025).

77 Njei, B. et al. Lean MASLD in U.S. Adults: Undiagnosed Burden and Metabolic Features from NHANES 2017-2020. J Gen Intern Med (2025).

78 Wang, Q. et al. Non-obese histologically confirmed NASH patients with abnormal liver biochemistry have more advanced fibrosis. Hepatol Int 13, 766–776 (2019).

79 The Diabetes Control and Complications Trial Research Group. The effect of intensive treatment of diabetes on the development and progression of long-term complications in insulin-dependent diabetes mellitus. N Engl J Med 329, 977–986 (1993).

80 UK Prospective Diabetes Study (UKPDS) Group. Intensive blood-glucose control with sulphonylureas or insulin compared with conventional treatment and risk of complications in patients with type 2 diabetes (UKPDS 33). Lancet 352, 837–853 (1998).

81 Chen, J. et al. Inhibition of O-GlcNAc transferase activates type I interferon-dependent antitumor immunity by bridging cGAS-STING pathway. Elife 13, RP94849 (2024).

82 Elvevold, K., Kyrrestad, I. & Smedsrød, B. Protocol for Isolation and Culture of Mouse Hepatocytes (HCs), Kupffer Cells (KCs), and Liver Sinusoidal Endothelial Cells (LSECs) in Analyses of Hepatic Drug Distribution. Methods Mol Biol 2434, 385–402 (2022).

83 Antwi, M. B. et al. Liver sinusoidal endothelial cells show reduced scavenger function and downregulation of Fc gamma receptor IIb, yet maintain a preserved fenestration in the Glmpgt/gt mouse model of slowly progressing liver fibrosis. PLoS One 18, e0293526 (2023).

84 Gnanachandran, K. et al. Sublethal stress from polypharmacy modulates scavenging function and fenestrations in mouse liver sinusoidal endothelial cells. J Pharmacol Exp Ther 393, 104955 (2026).

85 Schmidt, U., Weigert, M., Broaddus, C. & Myers, G. Cell Detection with Star-Convex Polygons. Medical Image Computing and Computer Assisted Intervention (MICCAI) 11071, 265–273 (2018).

86 Szafranska, K. et al. Quantitative analysis methods for studying fenestrations in liver sinusoidal endothelial cells. A comparative study. Micron 150, 103121 (2021).

87 Klaesson, A. et al. Improved efficiency of in situ protein analysis by proximity ligation using UnFold probes. Sci Rep 8, 5400 (2018).

88 Hegazy, M. et al. Proximity Ligation Assay for Detecting Protein-Protein Interactions and Protein Modifications in Cells and Tissues in Situ. Curr Protoc Cell Biol 89, e115 (2020).

89 Kojima, H. et al. Detection and imaging of nitric oxide with novel fluorescent indicators: diaminofluoresceins. Anal Chem 70, 2446–53 (1998).

90 Schindelin, J. et al. Fiji: an open-source platform for biological-image analysis. Nat Methods 9, 676–82 (2012).

