## Supplementary Information for "The hepatic sinusoid is a site of microvascular complications of hyperglycemia"

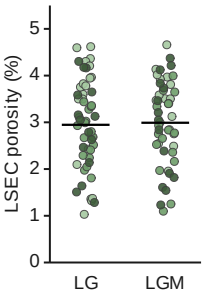

**Supplementary Fig. 1 | Mannitol does not alter LSEC porosity.** LSEC porosity in low glucose (5.5 mM) with and without mannitol (24.5 mM). Each dot is one cell, shaded by preparation. Horizontal line, mean of preparation means. n = 3 independent preparations. Two-tailed unpaired t-test.

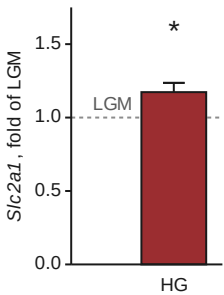

**Supplementary Fig. 2 | High glucose does not reduce *Slc2a1* expression.** *Slc2a1* transcript in HG relative to LGM. Bar, geometric mean. n = 5 independent preparations. Error bars, s.e.m. on the log2 scale. One-sample t-test on log2 ratios. \*P < 0.05 versus LGM.

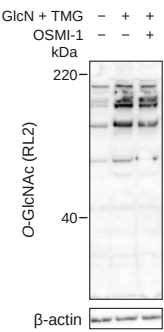

**Supplementary Fig. 3 | Glucosamine and Thiamet-G increase protein O-GlcNAcylation in the human LSEC.** Representative O-GlcNAc (RL2) immunoblot of primary human LSECs in low glucose, with GlcN + TMG, and with GlcN + TMG and OSMI-1. β-actin, loading control. n = 2 independent preparations.

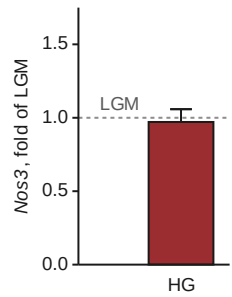

**Supplementary Fig. 4 | High glucose does not change *Nos3* expression.** *Nos3* transcript in HG relative to LGM. n = 5 independent preparations. Error bars, s.e.m. One-sample t-test on log2 ratios.

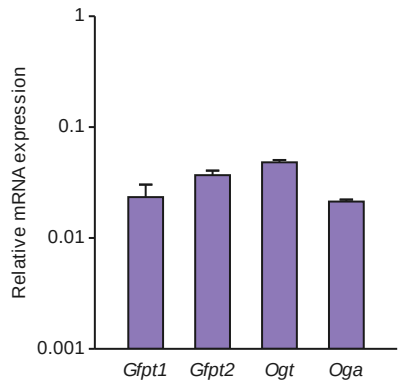

**Supplementary Fig. 5 | Expression of GFAT and the O-GlcNAc cycling enzymes in the LSEC.** Transcripts relative to reference genes, on the same scale as Fig. 4b and 4c. n = 20 samples. Error bars, s.e.m.

**Supplementary Table 1 | Metabolites after heptelidic acid.** Fold change, P and Benjamini-Hochberg q against control for each metabolite (n = 5 per arm). UDP-Hex, UDP-HexNAc, HexNAc and FBP are unresolved isomers, reported combined. Position is relative to the GAPDH step of glycolysis. Detected is the percentage of all samples in which the compound was above the detection limit.

| Figure label | Metabolon compound name | Position | Fold | P | q | Detected (%) |
| --- | --- | --- | --- | --- | --- | --- |
| G6P | glucose 6-phosphate | above | 41.51 | <0.000001 | <0.0001 | 17.9 |
| Galactose 6-P | galactose 6-phosphate | side branch | 2.62 | 0.000001 | 0.00004 | 14.3 |
| UDP-Hex | UDP-glucose/UDP-galactose | branch | 2.25 | 0.0001 | 0.0015 | 25.0 |
| 3PG | 3-phosphoglycerate | below | 2.13 | 0.0669 | 0.2631 | 50.0 |
| UDP-HexNAc | UDP-N-acetylglucosamine/galactosamine | branch | 2.01 | 0.0007 | 0.0071 | 21.4 |
| HexNAc | N-acetylglucosamine/N-acetylgalactosamine | salvage input | 1.53 | 0.0368 | 0.1824 | 92.9 |
| FBP | fructose 1,6-diphosphate/glucose 1,6-diphosphate/myo-inositol diphosphates | above | 1.30 | 0.2780 | 0.5969 | 92.9 |
| Glycerate | glycerate | below | 0.37 | 0.1382 | 0.4002 | 89.3 |
| Lactate | lactate | below | 0.14 | 0.000029 | 0.0005 | 100.0 |
| Glucose | glucose | above | 0.05 | 0.2169 | 0.5334 | 100.0 |

**Supplementary Table 2 | TaqMan gene expression assays.** Assay identifiers for every transcript measured by qRT-PCR.

| Gene symbol | Assay ID |
| --- | --- |
| *LSEC identity and capillarization* |  |
| *Stab1* | Mm00460390_m1 |
| *Stab2* | Mm00454684_m1 |
| *Fcgr2b* | Mm00438875_m1 |
| *Cd34* | Mm00519283_m1 |
| *Pecam1* | Mm01242576_m1 |
| *Glucose transporters* |  |
| *Slc2a1* | Mm00441480_m1 |
| *Slc2a2* | Mm00446229_m1 |
| *Slc2a3* | Mm00441483_m1 |
| *Slc2a4* | Mm00436615_m1 |
| *Hexosamine salvage and synthesis* |  |
| *Nagk* | Mm00479878_m1 |
| *Amdhd2* | Mm01302556_m1 |
| *Gnpda1* | Mm01254912_gH |
| *Gnpda2* | Mm00503573_m1 |
| *Pgm3* | Mm01144498_m1 |
| *Uap1* | Mm01281909_m1 |
| *Reference genes* |  |
| *B2m* | Mm00437762_m1 |
| *Hprt* | Mm03024075_m1 |
| *Ppia* | Mm02342430_g1 |
| *Rpl13a* | Mm05910660_g1 |
